# CELL CYCLE STATE PREDICTION USING GRAPH NEURAL NETWORKS

**DOI:** 10.1101/2024.01.30.577893

**Authors:** Sayan Acharya, Aditya Ganguly, Ram Sarkar, Abin Jose

## Abstract

Mitosis is a crucial process ensuring the faithful transmission of the genetic information stored in the cell nucleus. Aberrations in this intricate process pose a significant threat to an organism’s health, leading to conditions like cancer and various diseases. Hence, the study of mitosis holds paramount importance. Recent investigations have involved manual and semiautomated analyses of time-lapse microscopy images to understand mitosis better. This paper introduces an approach for predicting mitosis stages, employing a Convolutional Neural Network (CNN) as the initial feature extractor, followed by a Graph Neural Network (GNN) for predicting cell cycle states. A distinctive timestamp is incorporated into the feature vectors, treating this information as a graph to leverage internal interactions for predicting the subsequent cell state. To assess performance, experiments were conducted on three datasets, demonstrating that our method exhibits comparable efficacy to state-of-the-art techniques.

## 1. INTRODUCTION

The study of cell state division is important because it provides insights into fundamental biological processes and has significant implications in fields like medicine and biotechnology [1]. Cell state division is crucial in understanding how diseases like cancer take root in the human body [2]. It is equally important for the development of drugs and regenerative medication as well [3]. It is useful in various sub-fields of biotechnology like cell culture study, fermentation, etc. [4]. Therefore, this ever-growing and medically relevant study of cell division was a huge motivational factor for this work.

### 1.1. Related work

With the advancement in the area of deep learning, now deep learning is extensively used for mitosis detection. Zhong et al., [5] proposed an unsupervised method for identifying the different stages of mitosis. They use features that are extracted from the time-lapse microscopy images of human tissue culture cells using CellCognition [6] software. Flow cytometry is another major advancement in the study of cell cycle [7]. However, this method lacks the temporal component of cell-cycle progression. As reported in [8, 9], DNA content could be used for identifying cell cycle phases and perfect segmentation of the cell nucleus is the major requirement for this method. For cell-cycle classification, Ferro et al. [10] proposed feature extraction from fluorescence images followed by K-means clustering to identify cell-cycle stages. Deep learning methods [11, 12, 13], to name a few, were used for feature extraction and classification. Jin et al. [14] proposed image classification based on WGAN-GP and ResNet, and they tried to address the data imbalance problem with the cell-cycle detection datasets. A deep neural network-based approach for classifying interphase cell-cycle stages in microscopy images was proposed by Narotamo et al. [15]. Detection of mitosis in fixed tissue samples using deep learning [16, 17, 18], was common, however, it was not extensively used in video sequences of cells undergoing mitosis. The similarity between images in different stages of mitosis will introduce high classification noise at the state transitions and authors showed that by introducing the time information into annotation this noise could be reduced. They used a supervised machine learning method with hidden Markov model [19]. Jose et al. [20, 21] proposed a recurrent neural network (RNN)-based method for cell cycle stage identification. This method uses the RNN model with Gated Recurrent Units (GRU) for classifying the cell-cycle stages.

### 1.2. This paper

In this paper, we propose a novel approach in which we employ a pre-trained neural network model as a feature extractor to construct a graph network using time-series cell images. Subsequently, we train a Graph Neural Network (GNN) model [22][to predict the immediate next state of the cell. This method focuses on identifying connections between different time steps in a cell division process and treats this information as a graph, leveraging internal interactions to predict the subsequent cell state. Unlike traditional neural networks, which excel in processing grid-like data, GNNs adeptly handle graph-structured information. Our experiments, conducted on images captured using three different microscopes, demonstrate a key advantage of the GNN-based method over RNN-based approaches in classifying the images into different classes such as pre-mitosis, mitosis, and post-mitosis.

The main advantages of using the GNN model are:

1. No vanishing gradient issue.
2. Faster batch processing due to the absence of recurrent connections.
3. Arbitrary temporal window sizes can be used due to the versatility of the graph model. Therefore according to the need, more context of past images can be added to make the predictions better.

Specifically, GNNs are not restricted to a particular graph type or structure. This implies that graphs with more nodes (e.g., longer image sequences) and more edges (e.g., increased correlation between images in the sequence) can be incorporated into the network without requiring retraining, thereby enhancing the model’s versatility. Furthermore, as the GNN learns to utilize the entire graph information, it autonomously discerns the significance of timestamps and the correlation between images at different timestamps. This paper is organized as follows: Section 2 elucidates the proposed approach, Section 3 discusses the experimental setup and results, and Section 4 outlines conclusions and future work.

## 2. PROPOSED APPROACH

### 2.1. Formation of feature vectors

#### Overview of the method

The proposed method is explained in the flowchart shown in Fig. 1. Initially, a sequence of images is taken. The average length of pre-mitosis states and mitosis states are only 7-15 timestamps long with most of them having 10 timestamps. Therefore, 10 is chosen as the suitable temporal window size. Then the images are passed through a feature extractor pretrained on ImageNet, a ResNet-18 model, which generates a feature vector, **f** ∈ *ℛ* ^*d*^. The output of this is a stack of feature vectors, each corresponding to one image from the sequence. Two more features, the timestamp of the image in the sequence and the state of the cell are appended to **f**. These 2 features added are for the input image window which means they use the cell state and timestamp information of the images of the input window and not any information from the ground truth. A graph is then created, where the nodes are **f** and the edges are the similarity (e.g.: Cosine similarity) between different feature vectors. Then this graph along with these feature vectors is passed as input to the GNN. Inside the GNN, the inputs are passed through graph convolutional layers which does message-passing operations between the nodes and aggregate the information of the neighborhood of each node into that node. This learns the higher-level features of the graph structure and helps with the overall prediction task. Then, the output of the last convolution layer is passed through a filtering layer to aggregate the features. Then this aggregated feature vector is flattened into a final vector that is passed into the multilayer perceptron to make the final prediction.

**Fig. 1:**
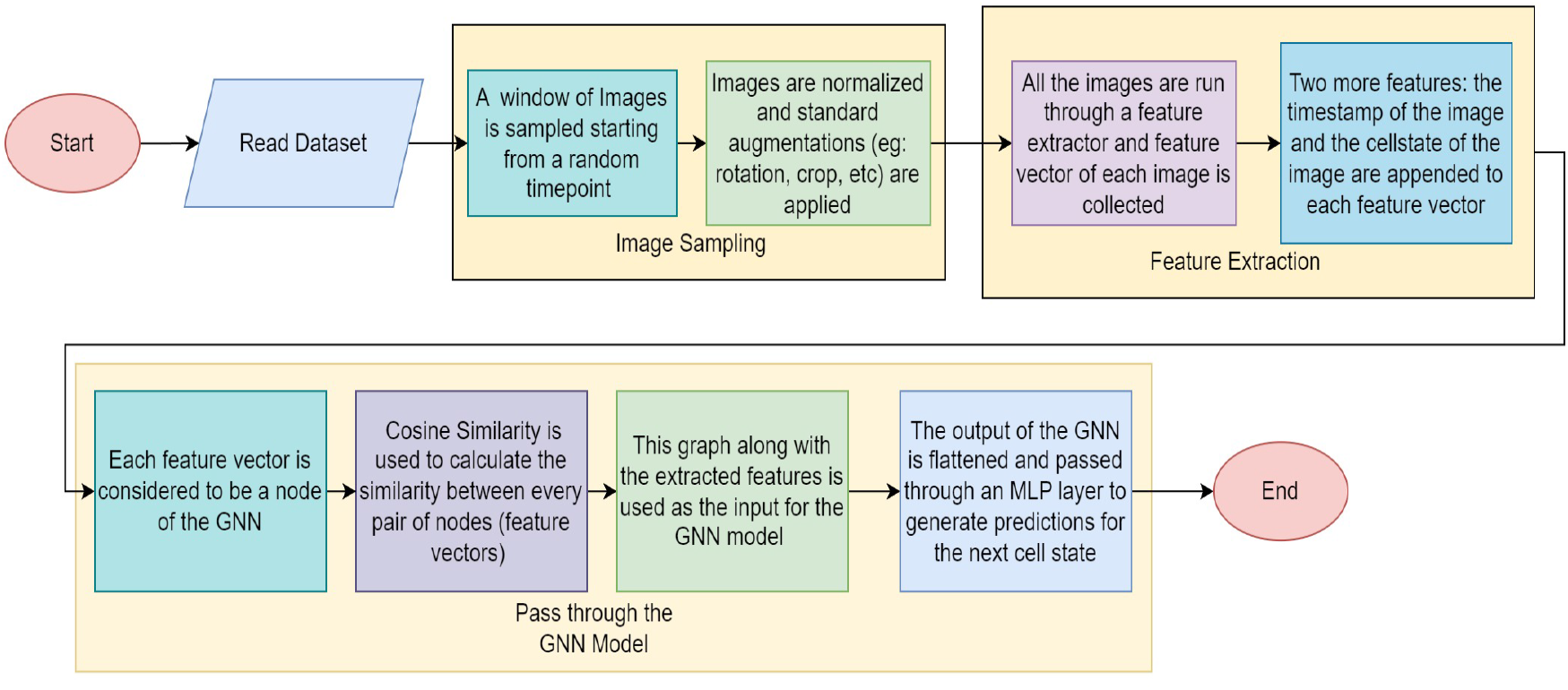
Flowchart of the pipeline of our proposed ResNet-18 based GNN model.

#### Preprocessing of the dataset

We are conducting our experiments on a 3-class dataset that contains time series image data of cell states during cell division. This is further elaborated upon in Section 3. The 3 cell-state dataset is mostly balanced with all the classes having a nearly equal number of images. These images are all in grayscale and in the shape of 96 *×* 96. So, these images are first rescaled to a size of 256 *×*256. We have used 256 *×*256 on the ResNet. Since we are not including the last dense layer, it is input size agnostic. Then to increase the robustness of the model, standard augmentation techniques like horizontal flipping, rotation, centre-crop, etc. are applied.

#### Sampling

For training, a contiguous sequence of *N* images is sampled from the dataset. Here this sequence will be passed as the input to the model whilst the state of the image just after the last image of this sequence will be the target state for the prediction.

#### Feature extraction

For the feature extraction, the ResNet-18 [24] is used as a baseline model. For our experiments, we have also considered baselines ResNet-50 [25] and VGG-16 [26].

In order to improve the separation between features of different classes, the center loss is applied [27]. Center loss is a technique used in deep learning to enhance the discriminative power of neural network models. It extends the standard classification objective, often a softmax cross-entropy loss, by introducing an additional term. This new term, the center loss, operates by calculating the Euclidean distance between the features (**f**_*i*_) and their respective class centers 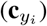, as 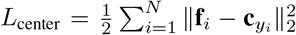 where, *N* is the number of images in the sequence, **f**_*i*_ is the feature vector of image *i, y*_*i*_ is the true class label of image *i*, and 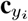 is the mean center of the features of true class of image *i*. These class centers are also trainable parameters. The aim is to minimize this distance, effectively encouraging features from the same class to cluster around their designated center. By incorporating the center loss in the overall training objective with an appropriate weighting factor (*λ*), the model is guided to learn a feature space where intraclass features are closer, contributing to improved feature discrimination.

#### 2.2 Prediction using GNN

For prediction using the GNN [23], the images are treated as the nodes of the graph, and the edges are defined as the similarity between the features of the images. Therefore, the graph structure is derived from the images themselves. At first, a random window is selected from the given dataset of sequential images. The task here is to predict the cell state of the next image from this window of images. This window of images is then passed to the feature extractor that gives the embedding of each image. Then a similarity function (e.g. Cosine similarity) is used to calculate edge values between every pair of nodes. This generates the windowSize *×* windowSize, adjacency matrix (**A**). At first, the adjacency matrix is normalized by dividing each edge weight with the geometric mean of the degrees of the nodes it is connected between. This gives the normalized Laplacian matrix to be used in the GNN. **D** is a diagonal matrix, where the diagonal elements are the summation of the weights of edges connected to the concerned node.

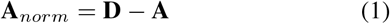

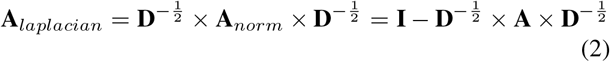

After normalization, this Laplacian matrix is passed through multiple graph convolutional layers to aggregate the neighborhood information. The temporal information is learned through the added feature in the feature vector. The succession of states and other inter-state information are learned from the graph structure of the window of images which incorporates the interaction of the states amongst one another. Then this matrix is passed through filtering layers to create an embedding of the information of the graph. The output of this layer is flattened out and passed to a multi-layer perceptron that produces the final prediction for what the next state would be. In this paper, a cross-entropy loss is used for back-propagation. The complete pipeline is shown in Fig. 2.

**Fig. 2:**
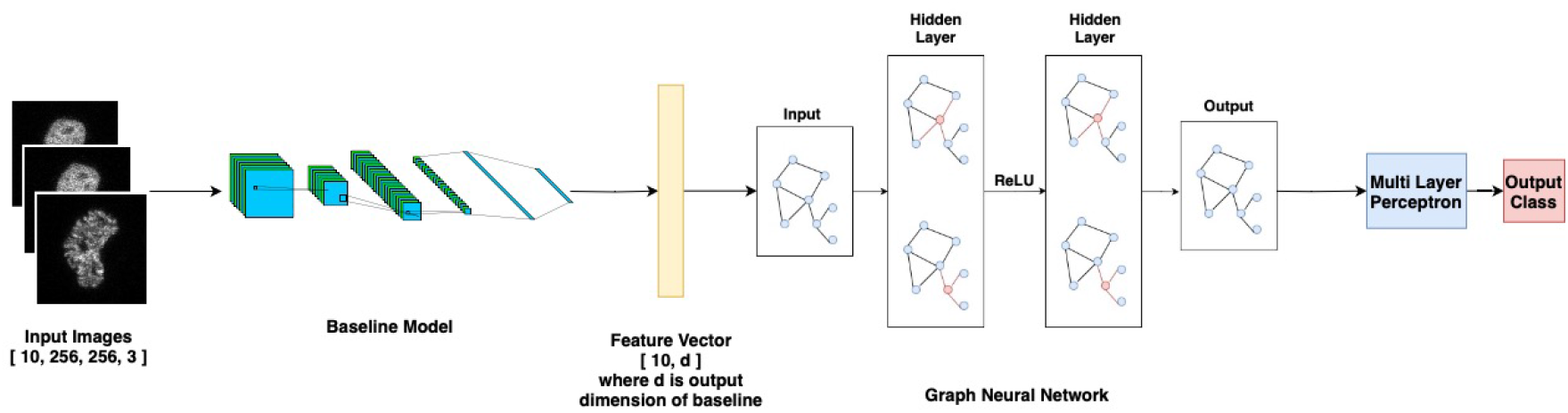
Architecture diagram of the end-to-end pipeline. The feature vector is passed through the GNN model, the output of the GNN is passed through a multi-layer perceptron (MLP) to predict the next cell cycle stage.

## 3. EXPERIMENTAL SETUP AND DATASETS

### The 3-class dataset

The 3-class dataset ^1^ contains time-series image data of cell states during cell division i.e., pre-mitosis (state 1), mitosis (state 2), and post-mitosis (state 3) as shown in Fig. 4. We have done the experiments ^2^ on three datasets provided by Moreno-Andres et al. [28]. These datasets contain microscopic image sequences of the mitotic process from human HeLa cells expressing H2B-mCherry as fluorescent chromatin marker. Extracted using LiveCellMiner [28], the datasets comprise microscopic image sequences depicting the mitosis process. Facilitating the analysis of mitotic phases in 2D+t microscopy images. Firstly, we have the LSD1 dataset, acquired using LSM5 live confocal microscope (Zeiss) and Zen software (Zeiss). Secondly, we have the LSM710 dataset where the cells are imaged using an LSM710 confocal microscope (Zeiss) and ZEN software (Zeiss). The third dataset is the NikonXLight dataset, the cells are imaged with the widefield module of a Ti2 Eclipse (Nikon) equipped with an LED light engine SpectraX and GFP/mCherry filter sets and using elements software (Nikon). These images of cells in various mitosis states can be seen in Fig. 3. In each time frame, the acquired images contain multiple cells. The software [28] actively tracks each cell’s position and subsequently extracts the tracked single-cell image. Cells not undergoing cell division are identified and excluded by LiveCellMiner. Note that for this dataset, the results are manually checked besides being filtered using LiveCellMiner.

**Fig. 3:**
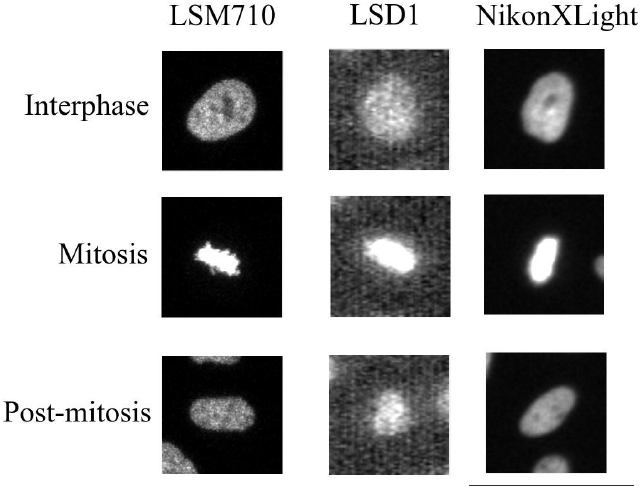
Cell images from the three datasets.

**Fig. 4:**
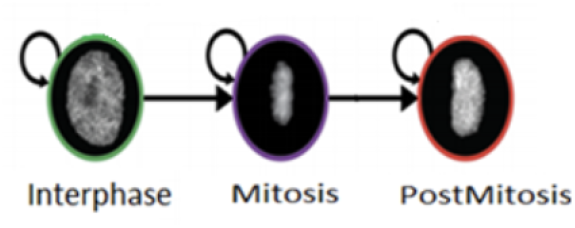
Transition diagram of three mitosis states.

A region is cropped around each tracked cell in each time frame if cell division occurs. Each of these cropped images maintains a resolution of 96×96 pixels, with the target cell positioned at the center. For each cell division sequence, 90 frames are available at this resolution. The dataset is categorized into three phases: interphase to early prophase as the first state, late prophase to anaphase as the next state, and telophase to interphase as the third state. The images, captured at a temporal resolution of three minutes, were utilized for training, incorporating a total of 1458 video sequences.

The cell state distribution in the images is mostly balanced with each class having almost the same number of images. The images are all grayscale images and have a dimension of 96 × 96. The images are already segmented such that the cell state of only one cell in the center is considered. As the images are time series data, the cell states of the images are automatically sorted in the numerical order of the states i.e. state 1 is followed by state 1 or state 2, state 2 is followed by state 2 or state 3, and state 3 is followed by state 3 or state 1. The images are accompanied by JSON files containing the timestamp of their occurrence.

### Training the baseline model

The images present are imbalanced with some classes having a higher count in a sequence and others having a lower count. As this would adversely affect training, for each sequence, counts are recorded for each class. During training time, the images from each class are sampled according to this distribution so that the number of images of each class in each batch is roughly the same. The model is trained for a total of 150 epochs. In each epoch, the model is trained on 50 batches. A batch size of 8 is used during the training process. The optimizer used is the Adam and the learning rate used is 0.00005.

### Training the Graph Neural Network

The images of the baseline model are used to create the adjacency matrix for the graph. A windowSize *×* windowSize matrix is populated where the entries denote similarity between the feature vector of each image of the window of images. Then this matrix along with the feature vectors is passed to the GNN and trained for 50 epochs. Each epoch has 25 batches. The optimizer used is Adam with an initial learning rate of 1e-3.

### Evaluation criteria and hyperparameters

The evaluation criteria for judging the models is the accuracy of the predicted next state. The loss is also monitored at every epoch to check the progression of the training as well. There are multiple hyperparameters in the entire pipeline which is finetuned using grid search approach [29], and they are explained below, with the respective values used in the experiments in brackets.

- *K*: This hyperparameter controls the number of times convolution is applied on the Graph Neural Network (6).
- *L*: This hyperparameter controls how many layers of the convolutional network are applied (8).
- *F* ^*′*^: This hyperparameter controls how many features the graph is boiled down to after the convolutional layers (64).
- HidSize: This hyperparameter controls the size of the hidden layers of the MLP layer (256).
- windowSize: This hyperparameter controls the window size of how many cell images to take into account for prediction (10).
- Lambda: This hyperparameter controls the weightage given to the center loss during the calculation of the total loss (1.0).

## 4. EXPERIMENTAL RESULTS

We have tried different backbones and for each baseline trained it with and without the inclusion of center loss during training. The accuracy shown is the accuracy obtained by the GNN on the prediction task when using the embedding of the feature extractor. For fairness of the evaluation, we kept a fixed GNN architecture whose size was determined by testing the performance on various datasets. In the filtering stage shown in Fig. 1, all the features derived from the previous GNN units are filtered, flattened, and passed to the MLP. Now the ResNet-50 has longer feature vector sizes (2048) whilst the VGG-16 and ResNet-18 have shorter feature vector sizes (512). Because of this, the GNN architecture should have been made more complex, but we did not do that and chose a size big enough (see the parameters mentioned in Section 3) to account for the complexities of all the feature extractors (having a bigger network tends to overfit a lot in the GNN due to the oversmoothing issue [30]).

Table 1 shows a set of results where we tested our whole GNN pipeline with different feature extractors on different cell tracking datasets. The ResNet-18 seems to be the best feature extractor for our pipeline.

**Table 1:** Performance of baseline models on 3 datasets.

| Feature vector | Dataset | Centreloss | No Centreloss |
| --- | --- | --- | --- |
| ResNet-50 | LSM710 | 96.34% | 95.05% |
| VGG-16 | LSM710 | 95.35% | **96.75%** |
| ResNet-18 | LSM710 | 95.65% | 95.50% |
| ResNet-50 | LSD1 | 96.50% | 96.45% |
| VGG-16 | LSD1 | 94.10% | 96.95% |
| ResNet-18 | LSD1 | **97.05%** | 94.95% |
| ResNet-50 | NikonXLight | 95.25 % | 95.20% |
| VGG-16 | NikonXLight | 93.10% | 93.05% |
| ResNet-18 | NikonXLight | **95.85%** | 93.90% |

We have also provided a side-by-side comparison with the frame-to-frame accuracy of the method in the paper [31]. However, the domain of the work is different. Here in this paper, we are predicting the state of the next image from a window of previous images whilst in the aforementioned paper, the authors have measured the individual classification accuracy of just the frame in question. Therefore, their method does not give a direct comparison with our method. The models in question are ResNet-18 in both cases and for our method, centreloss is considered. The comparison is provided in the Table 2.

**Table 2:** Accuracy comparison with RNN-based method [31].

| Dataset | Theirs | Ours |
| --- | --- | --- |
| LSM710 | 99.565 ± 0.040% | 95.65% |
| LSD1 | 99.572 ± 0.030% | 97.05% |
| NikonXLight | 99.292 ± 0.058 % | 95.85% |

The confusion matrices for the ResNet-18 baseline are provided for each of the three datasets in Fig 5. Most of the error cases are between adjoining stages in the time series, that is states 1,2, or states 2,3. This is because the data is sequential, as the cell moves from states 1 to 2 and then 3 the images at the boundary of these states cause prediction inaccuracies for the model. The t-SNE plot in 2D for a sample of features from the test set obtained from the ResNet-18 baseline models when using center loss is shown in Fig. 6. The mitosis class is clearly forming a separate sub-cluster for all three datasets. This is the expected behavior as the mitosis class frames are different from the post-mitosis and pre-mitosis frames. Furthermore, the cells after going through mitosis grow back to the original pre-mitosis state and hence the feature space is expected to have this overlap. An interesting thing to notice here is that the models that produce well-centered feature vectors have better inter-class separation using the Cosine similarity. Therefore, those feature vectors produce better edge weights when it comes to GNN which in turn produces better results. It could explain the performance difference between ResNet models and VGG-16. The feature vectors produced by the ResNet models are well-centered, but the ones produced by VGG-16 are such that the inter-class angular separation is low.

**Fig. 5:**
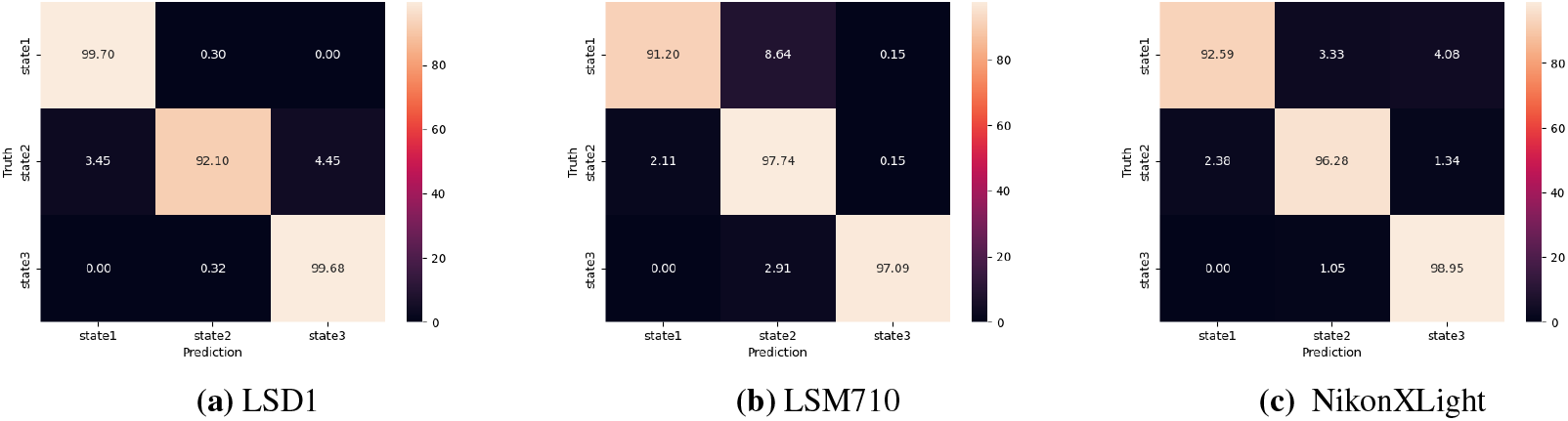
Confusion matrices for different datasets trained with ResNet-18 model.

**Fig. 6:**
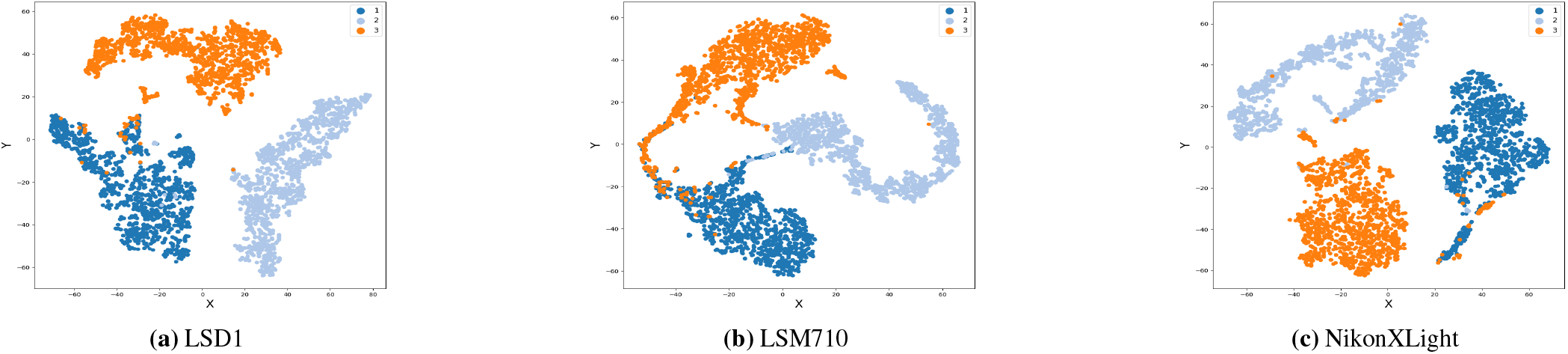
t-SNE plots for different datasets using ResNet-18 baseline. The overlap between pre-mitosis (1) and post-mitosis (3) states is expected as the frames have similar image content during these two states. The mitosis class (2) forms a separate sub-cluster.

### Error cases

As the cell division progresses, it cycles between the stages. When the division occurs, the cell goes through a lot of changes i.e. aggregation, and aligning of chromosomes; splitting of chromatids, de-condensation of chromosomes, and formation of the nucleus [32]. Due to so many different changes that a cell goes through during mitosis, it is fairly easy to tell it apart from the other 2 states. However, the difference between pre-mitosis and post-mitosis is harder to tell. Therefore in some cases, the proposed model becomes confused between these 2 states and creates the wrong type of feature vectors leading to wrong prediction. This is also evident from the t-SNE plots.

### Limitations

Although this method is novel and works well in most cases, some factors can limit the performance of our model and are discussed here: 1. Class imbalance: This model is very dependent on the amount of data received per class. Therefore, if the distribution of the classes is not uniform then the performance of this model is affected. 2. Dependency on feature vectors: As this model uses a Cosine similarity for the feature vectors, when the feature vectors have a greater angular separation between them, the similarity value should be higher. Now if it is the case that, the feature vectors have a worse angular separation then, the network produced by the Cosine similarity will not be very distinctive which will hinder the performance of the network. Therefore if the feature vectors are closely located at a particular spot away from the origin then their Cosine similarity will be very high. Therefore, the edges constructed tend to have similar values and do not convey much useful information. This ends up leading to a drop in performance and slow convergence of the loss.

## 5. CONCLUSIONS AND FUTURE WORK

In this paper, we have investigated the task of predicting the subsequent cell state by analyzing a sequence of cell images along with their respective states leading up to the prediction timestamp. Our approach involves utilizing a feature extractor based on CNN and GNN models. Various CNN models are employed to extract feature vectors from the given cell images, and these vectors form the basis of a graph network. The GNN is then trained using this graph network to forecast the next cell state. An important aspect of this method is its incorporation of GNN, and it remains independent of the graph size. Consequently, the parameter ’windowSize’ can be adjusted as needed, while maintaining the overall structure. Furthermore, the model considers interactions among all states leading to the prediction, resulting in a highly robust performance across a diverse range of feature extractor backbones and yielding satisfactory outcomes. Given the dynamic nature of advancements in the study of cellular mechanisms, there are ample opportunities for further exploration using deep learning techniques in the domain of cell state division. Future research directions could include exploring various feature extraction methods employing transformers, extending the analysis to 6-class state datasets, and other avenues for better predictive models.

## Footnotes

1 Dataset Repository: https://osf.io/8qdgm/

2 Code: https://github.com/SayanAcharya2002/CellTrackingPaper

## Notes

https://osf.io/8qdgm/

### Competing Interest Statement

The authors have declared no competing interest.

